# In vitro activity of cefiderocol against carbapenemase-producing and meropenem-nonsusceptible gram-negative bacteria collected in the Japan Antimicrobial Resistant Bacterial Surveillance (JARBS-GNR)

**DOI:** 10.1101/2024.02.14.580343

**Authors:** Shizuo Kayama, Sayoko Kawakami, Kohei Kondo, Norikazu Kitamura, Liansheng Yu, Wataru Hayashi, Koji Yahara, Yo Sugawara, Motoyuki Sugai

## Abstract

The treatment options available for infections caused by multidrug-resistant gram-negative pathogens are often limited. Cefiderocol (CFDC) is a novel siderophore cephalosporin that exhibits activity against multidrug-resistant gram-negative pathogens. Several studies have reported the in vitro activity of CFDC using clinical isolates from Europe, the United States, and China. However, no large-scale studies on the in vitro activities of CFDC have been conducted using all isolates with available genomic backgrounds based on whole-genome sequencing (WGS). We evaluated the antimicrobial activities of CFDC, ceftolozane/tazobactam (CTLZ/TAZ), imipenem-relebactam (IPM/REL), and ceftazidime/avibactam (CAZ/AVI) against carbapenemase-producing Enterobacterales, carbapenemase-non-producing meropenem-nonsusceptible Enterobacterales, and carbapenemase-producing non-fermentative bacteria. We selected 603 isolates (528 Enterobacterales, 18 *Pseudomonas aeruginosa*, and 57 *Acinetobacter* spp.) from the recent surveillance of clinical isolates in Japan using WGS data. Among these, 97.7% (300/307 strains) of carbapenemase-producing Enterobacterales, 100% (18/18 strains) of carbapenemase-producing *P. aeruginosa*, and 91.2% (52/57 strains) of carbapenemase-producing *Acinetobacter* spp. were susceptible to CFDC, showing better antimicrobial activity than the other antimicrobial agents evaluated in this study. In addition, CFDC was highly effective against class A, B, and D β-lactamase harboring isolates when compared to the other antimicrobial agents in this study. While β-lactam antibiotics were essentially ineffective against CFDC-resistant Enterobacterales, minocycline was the most effective, and gentamicin and amikacin were also effective. This is the first large-scale study to systematically demonstrate the efficacy of CFDC using carbapenemase-producing strains with transparent genomic backgrounds.

## Introduction

Multidrug-resistant gram-negative bacteria are becoming increasingly prevalent worldwide, resulting in nosocomial infections that are difficult to treat. Third-generation cephalosporin (3GC)-resistant and carbapenem-resistant Enterobacterales, and carbapenem-resistant *Acinetobacter baumannii* and *Pseudomonas aeruginosa* are included in the World Health Organization priority list (1). Effective treatments for these bacteria are currently extremely limited; therefore, the development of new antibiotic agents, especially those with different mechanisms of action, is urgently required.

There is an increasing demand for new antibacterial agents to treat infections caused by multidrug-resistant bacteria. The antimicrobial agents under clinical development are mainly derived from well-established antimicrobial agents. Therefore, preexisting cross-resistance limits the activity of new agents against multidrug-resistant gram-negative pathogens (2). In addition, current investments in the development of new antimicrobial agents are insufficient, particularly in the field of synthetic low-molecular-weight compounds derived from natural products (3).

In recent years, new antibiotics, such as ceftolozane-tazobactam (CTLZ/TAZ), imipenem-relebactam (IPM/REL), and ceftazidime-avibactam (CAZ/AVI), have been approved for clinical use (4). Regarding CTLZ/TAZ, ceftolozane is a 3’-aminopyrazolium cephalosporin that is stable against overexpression of AmpC and shows remarkable activity against *P. aeruginosa* (5) and extended-spectrum β-lactamase (ESBL)-producing isolates but not carbapenemase-producing isolates (6). Regarding IPM/REL, relebactam inhibits class A and C but not class B or D β-lactamases (7). IPM/REL shows activity against most *Klebsiella pneumoniae* carbapenemase (KPC)-producing and carbapenem-resistant isolates of *P. aeruginosa* but not against those of *A. baumannii* (7, 8). Regarding CAZ/AVI, avibactam binds to class A and C β-lactamases and certain oxacillinases (OXAs). Therefore, CAZ/AVI is highly active against KPC-producing isolates and has become the first-line therapy for these infections (4). However, avibactam does not inhibit metallo-β-lactamases (MBLs) such as New Delhi metallo-β-lactamases (NDMs) (9). Theoretically, these three new antimicrobial agents are unsuitable for the imipenemase (IMP)-type β-lactamases (class B MBL), which are dominant in Japan (10). Therefore, there remain concerns regarding their use as an essential option for antimicrobial agents.

Cefiderocol (CFDC), reported as a novel siderophore cephalosporin conjugated with a catechol moiety (11), shows potent static activity against carbapenem-resistant Enterobacterales, *P. aeruginosa*, and *A. baumannii* (12–14). A previous study revealed that the spectrum of CFDC includes a wide range of carbapenemase-harboring isolates, including serine (KPC, OXA) and MBLs (IMP, NDM, and Verona integron-encoded metallo-β-lactamase (VIM)) (14–16), and excellent stability of CFDC against hydrolysis by a variety of β-lactamases, including classes A, B, C, and D (4). However, studies have not yet been conducted in Japan, where the IMP-type MBL is dominant, and there have been no large-scale studies on the efficacy of CFDC against these isolates; all of these genomic backgrounds are available. For example, the resistance mechanisms to CFDC of *E. coli* have been well investigated, and previous reports (17, 18) have identified that CFDC resistance in *E. coli* is caused by a combination of 1) truncation of the colicin receptor CirA, 2) penicillin-binding protein 3 (PBP3) mutation caused by YRIN(K) insertion, and 3) the presence of *bla*_NDM_ (encoding the MBL NDM-1). The WGS data obtained in this study allowed a more detailed investigation of the mechanisms of resistance to CFDC.

Previously, we conducted national antimicrobial resistance genomic surveillance (Japan Antimicrobial Resistance Bacterial Surveillance (JARBS)) (19). All collected isolates in this surveillance (25,043 strains) were initially screened by performing a multiplex PCR assay for ESBL and carbapenemase genes. Selected strains were then subjected to genome sequencing and standardized quantitative antimicrobial susceptibility testing. These strains consisted of 4,195 isolates of *E. coli* and *K. pneumoniae* resistant to 3GCs and Enterobacterales (20), 671 isolates of *P. aeruginosa*, and 64 isolates of *Acinetobacter* spp.

This study aimed to investigate the in vitro activities of CFDC, CTLZ/TAZ, IPM/REL, CAZ/AVI, and carbapenems against Enterobacterales, *P. aeruginosa*, and *A. baumannii* collected during the national antimicrobial resistance genomic surveillance, JARBS (19), where IMP-type MBLs are dominant, unlike those in Europe and the United States.

## Materials and methods

### Bacterial isolates

*E. coli* and *K. pneumoniae* isolates resistant to 3GCs, Enterobacterales with reduced susceptibility to meropenem (MEPM), *P. aeruginosa*, and *Acinetobacter* spp. were collected in 2019 and 2020 from 175 hospitals through the JARBS. Paired-end sequencing (2×150 bp) of the isolates was performed as previously described (20). Of these, 528 Enterobacterales, 18 carbapenemase-producing *P. aeruginosa,* and 57 *Acinetobacter* spp. were used in this study. A breakdown of the 528 isolates examined is shown in Fig. 1.

**FIG 1.**
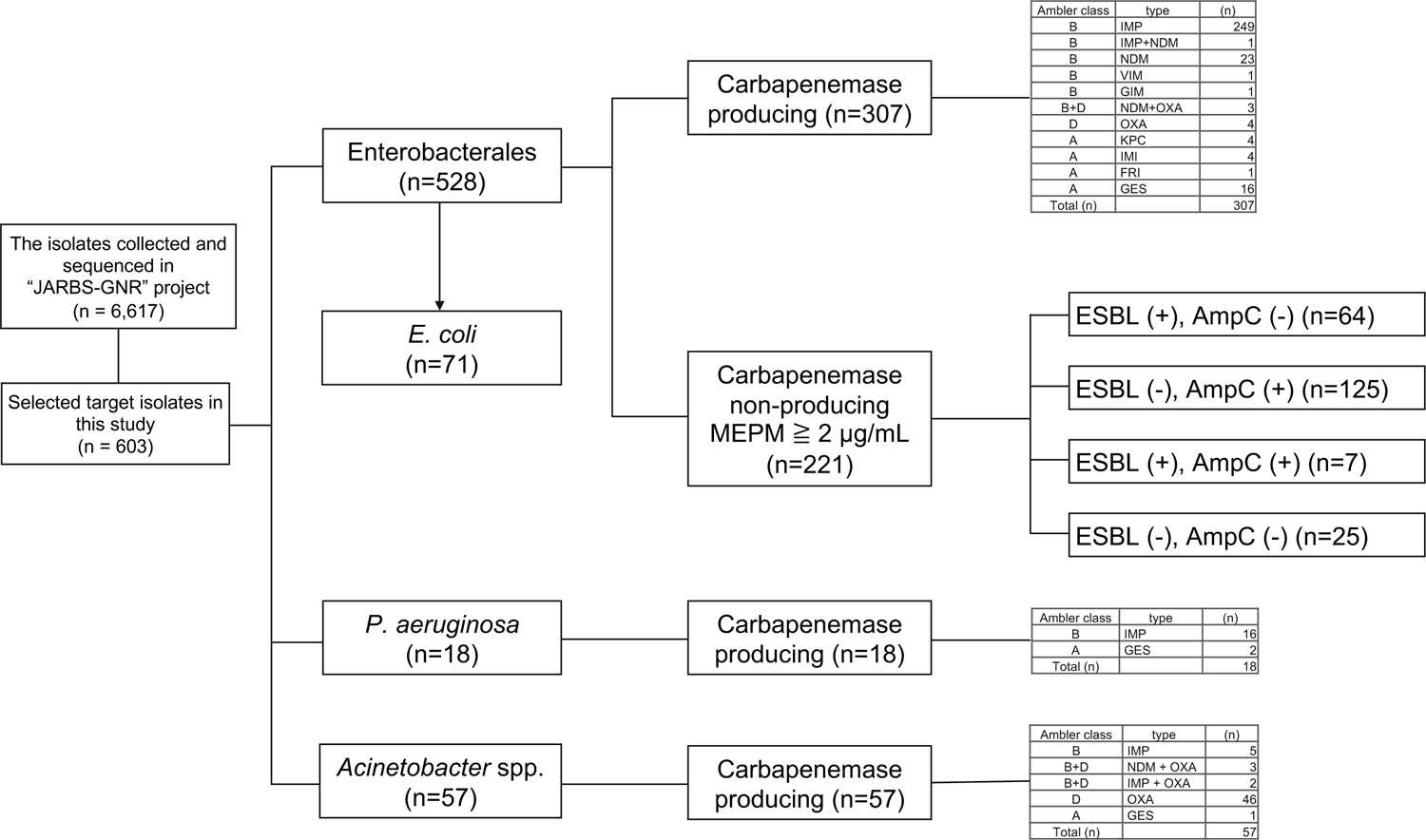
Summary of the 603 isolates used in this study.

### Antimicrobial susceptibility testing

CFDC, CTLZ/TAZ, IPM/REL, and CAZ/AVI were prepared at the research laboratories of Shionogi & Co., Ltd. (Osaka, Japan) as previously described (11). MICs were determined by broth microdilution, according to the protocol of the Clinical and Laboratory Standards Institute (CLSI 2022, 32nd Edition; Document M100). To determine the MIC of CFDC, iron-depleted cation-adjusted Mueller-Hinton broth was prepared as previously described (11). The MICs for the following antibacterial agents were determined using the broth microdilution method implemented in a MicroScan WalkAway (Beckman Coulter) using the NEG MIC 3.31E or NEG MIC NF 1 J panels (Beckman Coulter): piperacillin/tazobactam (PIPC/TAZ), cefotaxime (CTX), ceftazidime (CAZ), cefepime (CFPM), imipenem (IPM), MEPM, doripenem (DRPM), gentamicin (GM), tobramycin (TOB), amikacin (AMK), levofloxacin (LVFX), ciprofloxacin (CPFX), minocycline (MINO), sulfamethoxazole/trimethoprim (ST), chloramphenicol (CP), and colistin (CL). The MIC cutoff values were used according to the CLSI 2022, 32nd Edition; Document M100. *E. coli* ATCC 25922 and *P. aeruginosa* ATCC 27853 were used as control isolates.

## Results

### Antimicrobial activity against carbapenemase-producing Enterobacterales

In this study, CFDC showed potent antimicrobial activity against 307 carbapenemase-producing Enterobacterales (CPEs) (Fig. 2A). In contrast, CAZ/AVI or CTLZ/TAZ showed low antimicrobial activity. Of the isolates, 31.3% showed an intermediate resistance to IPM/REL, a similar susceptibility pattern to IPM. The rates of susceptibility of the 307 CPEs according to the type of carbapenemase (IMP, NDM, KPC, OXA, VIM, IMI, FRI, GES, and GIM) are listed in Supplemental Fig. 1. CFDC revealed high potency against isolates producing class A, B, and D β-lactamases. However, some isolates harboring NDMs were slightly more resistant to CFDC than those carrying the other β-lactamases (Fig. 2B). In addition, isolates producing class B β-lactamases were less susceptible to CAZ/AVI, IPM/REL, and CTLZ/TAZ. CAZ/AVI showed a significant effect against class A and D β-lactamase harboring isolates, whereas IPM/REL and CTLZ/TAZ showed limited efficacy. The MIC_50_ and MIC_90_ of CFDC against all CPEs were 0.25 mg/ml and 2 mg/ml, respectively (Supplemental Fig. 1). Other tested antibiotic agents, including CTLZ/TAZ, IPM/REL, and CAZ/AVI, were less active than CFDC against most carbapenemase producers, with an MIC_90_ of ≧64 mg/ml (CTLZ/TAZ), 16 mg/ml (IPM/REL), and ≧64 mg/ml (CAZ/AVI).

**FIG 2.**
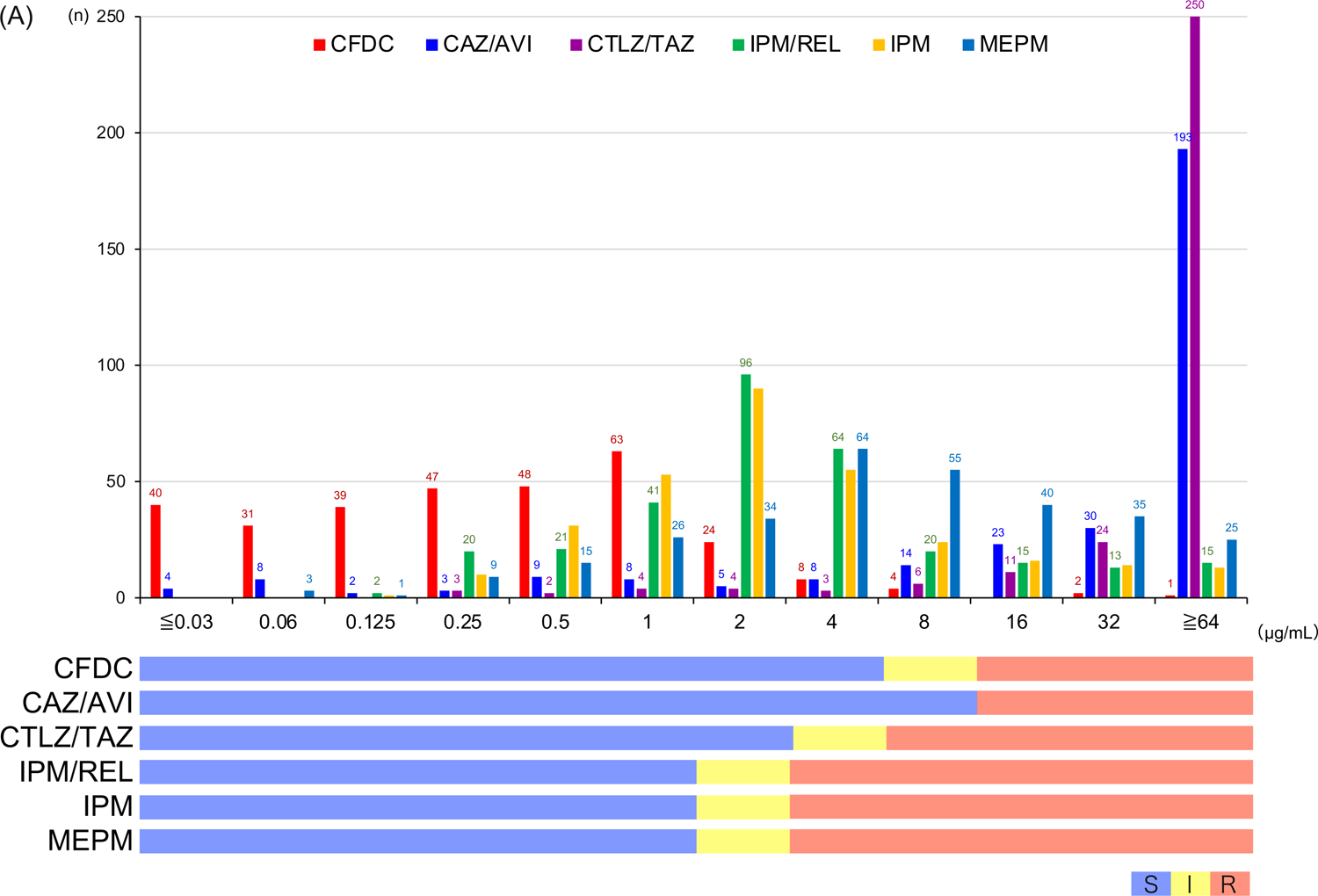
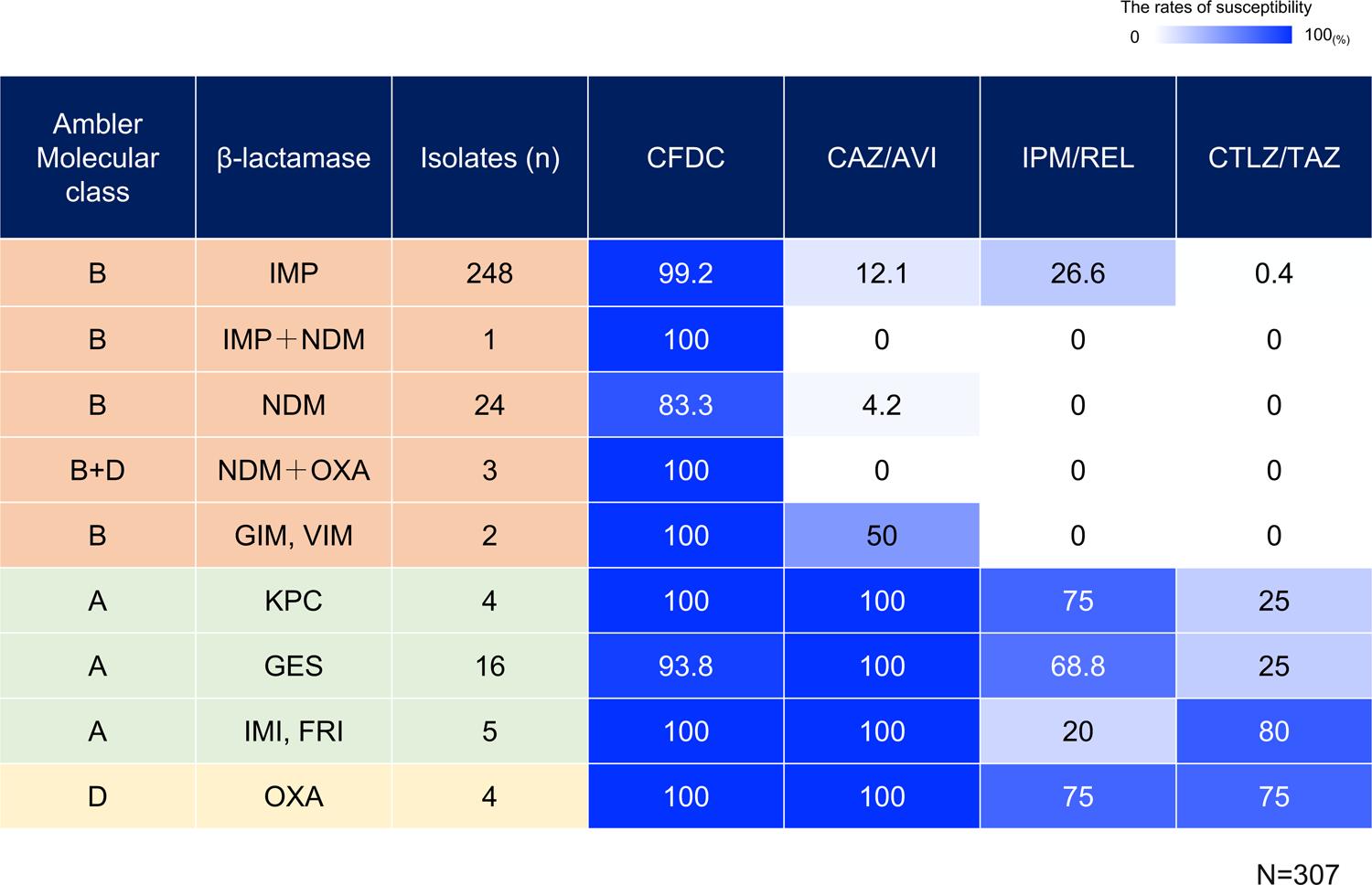
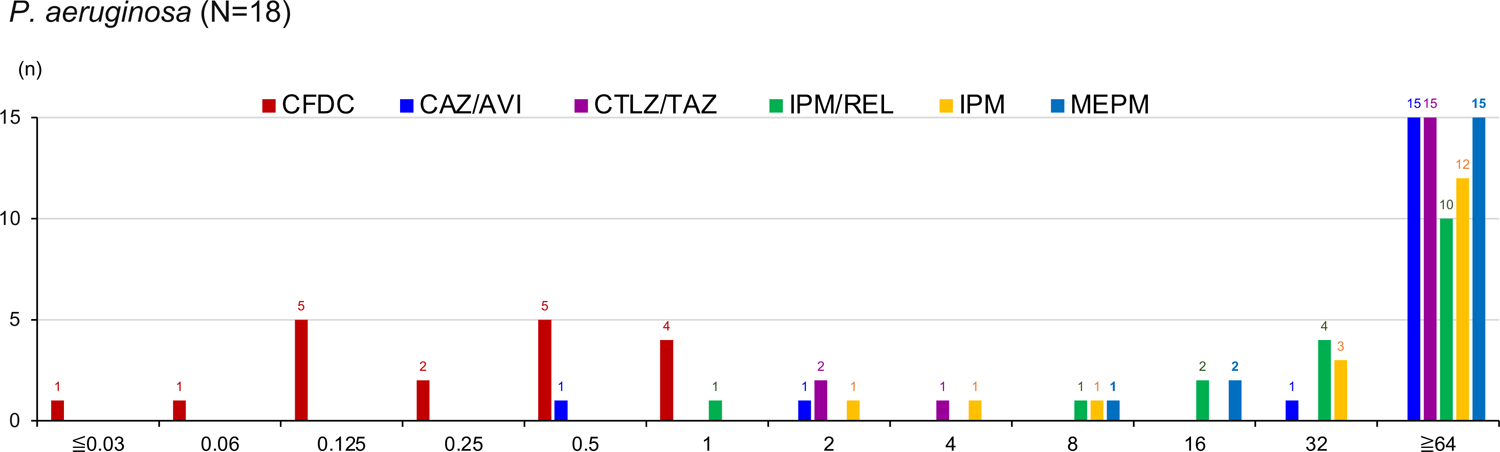
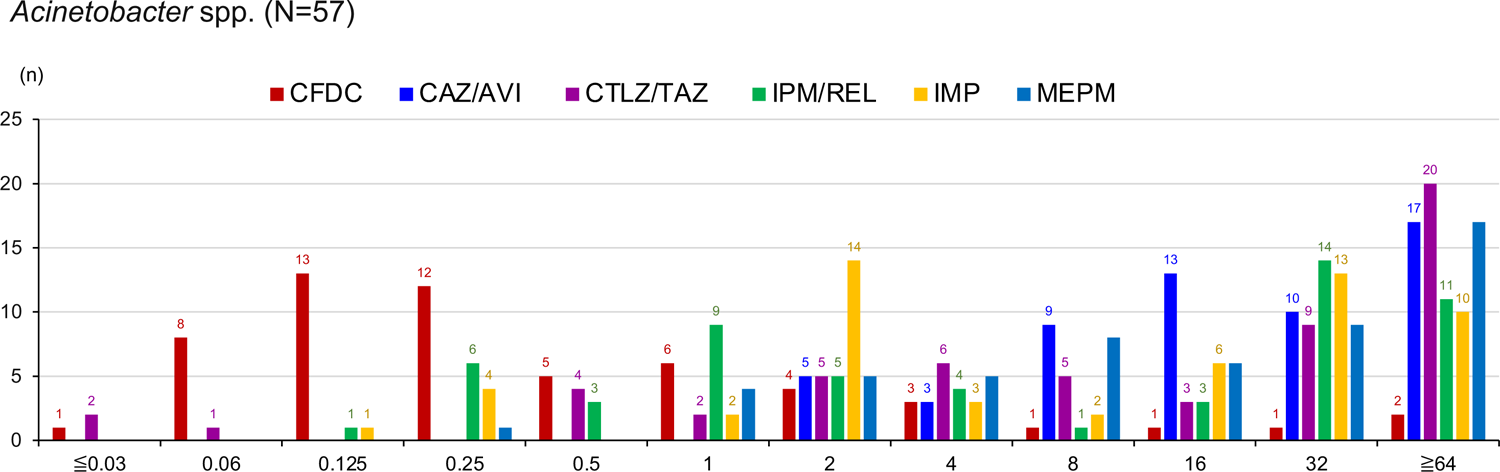
Summary of the MIC values in this study. (**A**) Bar chart of the distribution of antibiotic susceptibility of the 307 CPEs. CFDC is shown in red, CAZ/AVI in blue, CTLZ/TAZ in purple, IPM/REL in green, IPM in orange, and MEPM in pale blue. At the bottom of the figure, the clinical criteria for each antibiotic agent is color-coded (blue for susceptible, yellow for intermediate resistance, and red for resistant). (**B**) The effect of each antibiotic agent against the CPEs according to β-lactamase type. (**C**) and (**D**) Bar charts showing the distribution of antibiotic susceptibility for (**C**) *P. aeruginosa* and (**D**) *Acinetobacter* spp. CFDC is shown in red, CAZ/AVI in blue, CTLZ/TAZ in purple, IPM/REL in green, IPM in orange, and MEPM in pale blue.

### Antimicrobial activity against carbapenemase-producing non-fermentative bacteria

Similar to Enterobacterales, CFDC showed significant antimicrobial activity against carbapenemase-producing non-fermentative bacteria (Fig. 2C, 2D). No *P. aeruginosa* isolates were resistant to CFDC, although the number of isolates tested was limited. In contrast, CAZ/AVI, CTLZ/TAZ, and IPM/REL showed weaker activities than CFDC.

### The effect of CFDC and comparison among other antibiotic agents

CFDC was highly effective against all tested isolates, including carbapenemase-, ESBL-, and AmpC-producing Enterobacterales, carbapenemase-producing *P. aeruginosa*, and carbapenemase-producing *Acinetobacter* spp. However, none of the other antibiotic agents were as effective as CFDC (Fig. 3A). Regarding the 528 Enterobacterales (Fig. 3B), the β-lactams, including PIPC/TAZ, MEPM, DRPM, and IPM had almost no effect on CFDC-resistant isolates. In contrast, MINO was the most effective against CFDC-resistant isolates, and GM and AMK were also relatively effective. One isolate that showed CFDC resistance (≧64 μg/mL) was a *bla*_NDM-9_-carrying *E. coli* susceptible only to MINO. In this study, five isolates showed resistance to CFDC (Table 1), and these isolates were all resistant to MEPM, DRPM, and IPM. However, two isolates showing resistance to CFDC at 16 μg/mL were susceptible to CAZ/AVI and IPM/REL, but these isolates did not possess carbapenemase; they were *Enterobacter hormaechei* subsp. steigerwaltii carrying *bla*_ACT-15_ and *Klebsiella aerogenes* carrying unclassified AmpC β-lactamase.

**FIG 3.**
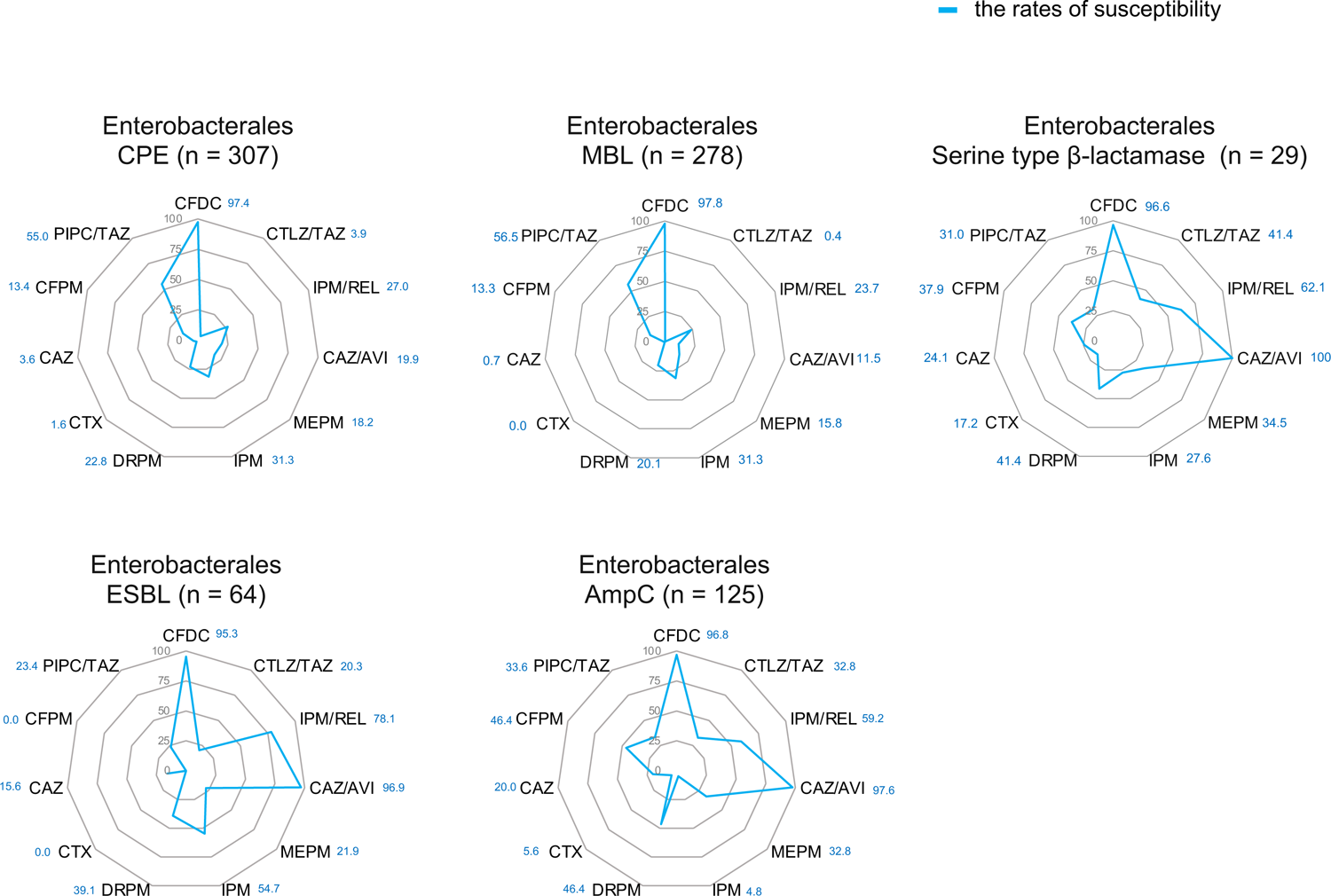
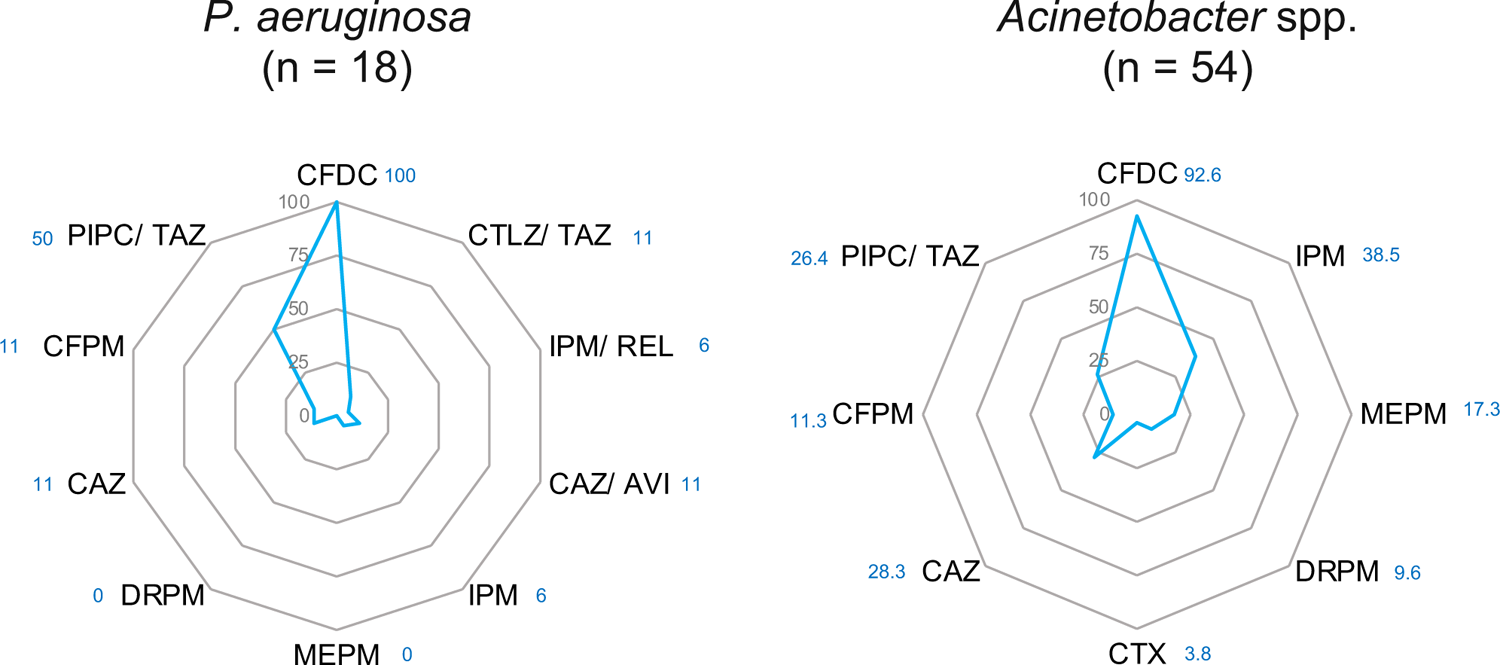
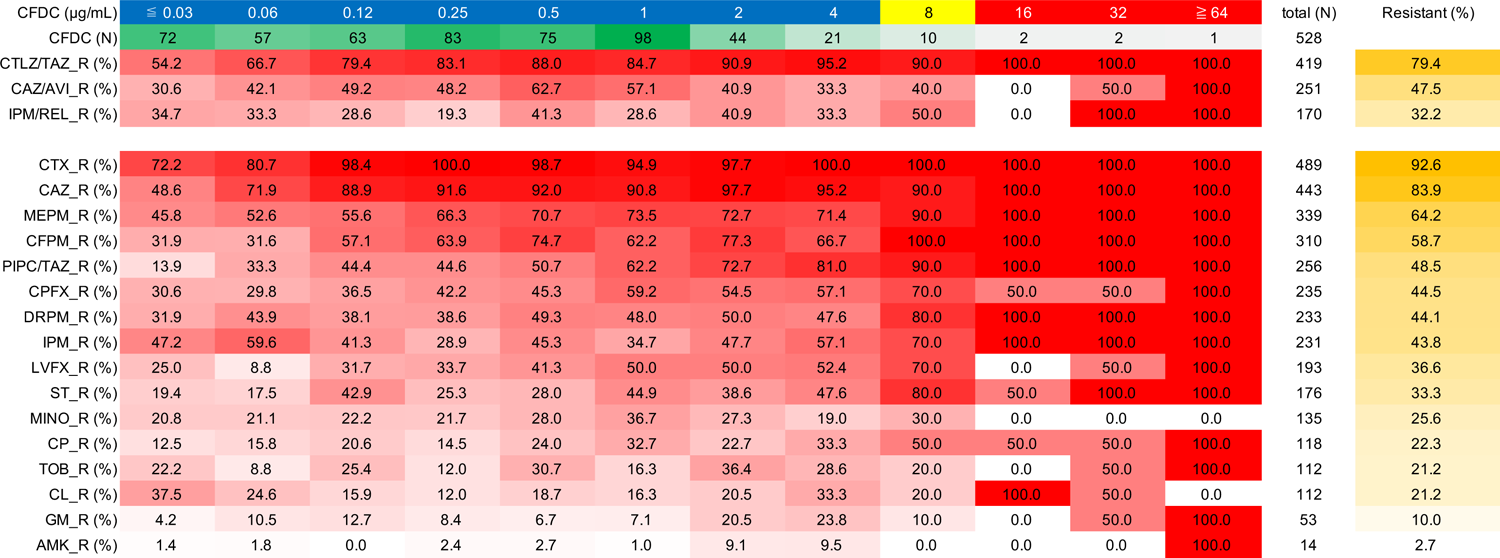
(**A**) Radar chart showing β-lactam susceptibility rates by bacterial species or resistance genes. Blue lines indicate percentages of susceptibility. (**B**) Relationship between CFDC MIC values and susceptibility to other antibiotic agents in 528 Enterobacterales isolates. The first row shows the concentrations of CFDC based on CLSI criteria (blue as susceptible, yellow as intermediate resistance, and red as resistant). The second row shows the number of isolates as a heatmap. The third and subsequent rows are in descending order according to the percentage of isolates resistant to each antibiotic agent. This table is divided into two parts by the horizontal white line. The upper part shows CAZ/AVI, CTLZ/TAZ, and IPM/REL (as new antibiotic agents), while the lower rows show the other typical antibiotic agents. The orange column on the right side shows the percentage of resistance to each antibiotic agent.

**Table 1.**
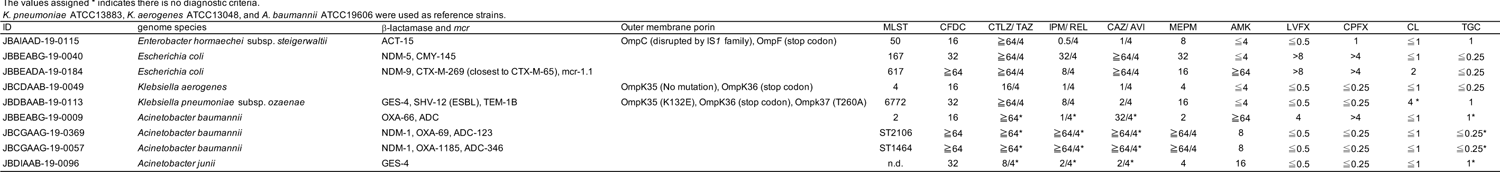
The resistant isolates against CFDC obtained in this study.

### Detailed characteristics of CFDC-resistant isolates obtained in this study

Among the isolates tested in this study, five Enterobacterales and four *Acinetobacter* spp. were resistant to CFDC (Table 1). Two *E. coli* isolates belonged to ST167 and ST617, whose sequence types were previously reported to be resistant to CFDC in China (17). Wang et al. (17) revealed the relationship between *bla*_NDM-5_ and CFDC resistance in *E. coli*. In the present study, one CFDC-resistant *E. coli* strain carried *bla*_NDM-9_ but not *bla*_NDM-5._ The CFDC-resistant *Klebsiella* and *Enterobacter* strains did not harbor NDM-type β-lactamase genes. In contrast, among 57 isolates, four *Acinetobacter* isolates showed resistance to CFDC, and all isolates with MIC values ≧64 mg/mL to CFDC possessed NDM-1.

### The mechanisms of CFDC-resistance in *E. coli*

In this study, 71 *E. coli* strains (Fig. 1) were analyzed for phenotypic (susceptibility to CFDC) and genotypic data. Among the 71 isolates, two were resistant, three showed intermediate resistance, and the remaining 66 were susceptible to CFDC. Among the two resistant isolates with MIC values >32 mg/mL, a combination of 1) the truncation of CirA, 2) PBP3 mutation caused by YRIN(K) insertion, and 3) the presence of *bla*_NDM_ was found (Fig. 4) (17, 18). Among the three isolates showing intermediate resistance to CFDC, two harbored *bla*_NDM_, two carried a CirA truncation, and one harbored a YRIN insertion in PBP3. In other words, intermediate resistant isolates possessed part of “the three CFDC-resistance factors” for acquiring CFDC resistance, resulting in intermediate resistance. Among the isolates that were susceptible to CFDC according to clinical diagnostic criteria but had low-susceptibility (MIC values between 1 and 4 µg/mL, the columns in Fig. 4 showing MIC values are shown in light blue), 13.6% (3/22) of isolates showed truncation of CirA (determined as <80% coverage), 50% (11/22) of isolates harbored a YRIN(K) insertion in PBP3, and 54.5% (12/22) of isolates carried *bla*_NDM_. None of the isolates possessed all three CFDC-resistance factors.

**FIG 4.**
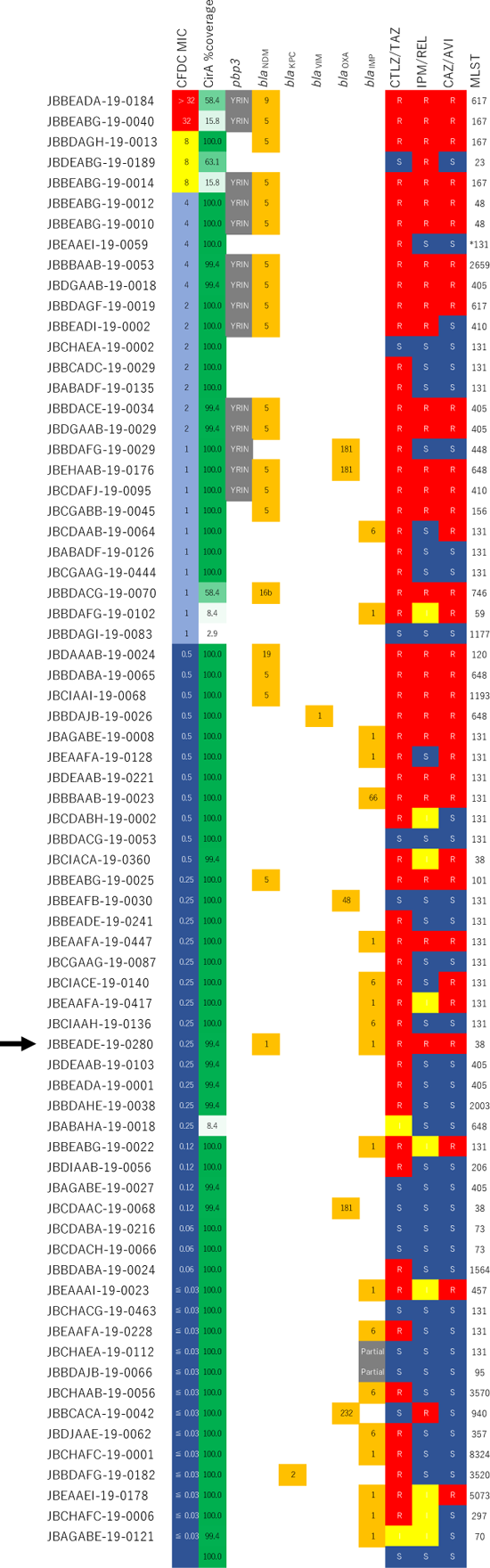
The genotype and susceptibility (CFDC and other antibiotic agents) of the 71 *E. coli* isolates in this study. The presence of the YRIN insertion in PBP3, along with the presence of carbapenemase genes, is indicated by color-coding in the columns for *pbp3*, *bla*_NDM_, *bla*_KPC_, *bla*_VIM_, *bla*_OXA_, and *bla*_IMP_. The numbers within the orange rectangles indicate the types of carbapenemases. The isolate harboring both NDM and IMP is indicated by the arrow.

Seven of the CFDC-low-susceptibility isolates did not possess any of these three CFDC-resistance factors. In contrast, the isolates that were highly susceptible to CFDC (column showing MIC values in dark blue, Fig. 4) possessed none of the three CFDC-resistant factors. Among the CFDC-susceptible isolates, five harbored NDM, but none possessed the other two CFDC-resistance factors.

## Discussion

CFDC is a novel siderophore cephalosporin conjugated to a catechol moiety (11). Previous studies have examined the susceptibility of numerous isolates to CFDC using only phenotypic information (21, 22). However, there have been no large-scale studies on CFDC using isolates with detailed genotypic information based on WGS. Furthermore, this study mainly focused on the effect of CFDC against bacteria carrying IMP-type β-lactamase, which is not prevalent in Europe and the United States.

Target isolates of Enterobacterales and non-fermentative bacteria were selected from 6,617 isolates from the JARBS project (Fig. 1). The most common CPE isolates harbored IMP-type β-lactamases (249 isolates), followed by the NDM-type (23 isolates). For Enterobacterales, we additionally selected 221 isolates that did not carry carbapenemases but showed reduced susceptibility to MEPM. These populations were further classified according to their β-lactamases (ESBL or *ampC*).

In this study, CFDC was effective (≦4 mg/ml) against 97.7% (300/307) of CPEs, 95.3% (61/64) of MEPM-nonsusceptible ESBL-producing Enterobacterales, 96.8% (121/125) of MEPM-nonsusceptible AmpC-producing Enterobacterales (Supplemental Fig. 1), 100% (18/18) of carbapenemase-producing *P. aeruginosa*, and 91.2% (52/57) of carbapenemase-producing *A. baumannii* (Supplemental Fig. 2). These results were mostly consistent with a previous report that CFDC MICs were 4 mg/ml for 97.0% of MEPM-nonsusceptible Enterobacterales (12).

None of the CFDC-resistant *Klebsiella* and *Enterobacter* harbored NDM-type β-lactamase genes, so the factors that add the resistance to CFDC against such species were presumed to differ from that of *E. coli*. The CFDC-resistant *Klebsiella* and *Enterobacter* harbored mutations in the outer membrane porin (Table 1); however, it is unclear whether these mutations are relevant to CFDC-resistance. Among the four CFDC-resistant *Acinetobacter* spp., one was carrying *piuA* involved in iron transfer disrupted by IS*Aba9* (23). Resistance mechanisms to CFDC in bacteria other than *E. coli* should be investigated further in the future.

Most of the 307 CPEs in this study were resistant to CAZ/AVI or CTLZ/TAZ (Fig. 2A). It has previously been reported that CAZ/AVI is mainly effective against class A, C, and some class D β-lactamase-producing isolates (9, 24, 25), and CTLZ/TAZ is effective against *P. aeruginosa* (26), ESBL (27), and AmpC-producers (28), but not against carbapenemase producers (6). The primary reason for the limited antimicrobial activity of CAZ/AVI and CTLZ/TAZ observed in this study was that these agents were ineffective against isolates carrying IMP-type MBLs (Fig. 2B). Regarding IPM/REL, intermediate resistant isolates were most common among the 307 CPEs and showed susceptibility patterns similar to those of IPM (Fig. 2A). This phenomenon was partially explained by the character of relebactam, which mainly inhibits class A and C β-lactamases but not class B MBL (29).

The isolates harboring multiple carbapenemases are included in the 307 CPEs in Fig. 2B. Among these isolates, IMP and NDM harboring *E. coli* (JBBEADE-19-0280) was susceptible to CFDC (MIC 0.25 µg/mL), and carried no obvious truncation of CirA or YRIN(K) insertion into PBP3 (indicated by the arrow, Fig. 4). There is a possibility that the presence of carbapenemases other than NDM, such as IMP or OXA, do not contribute to resistance to CFDC. Conversely, GES harboring isolates showed a slightly lower susceptibility rate (93.8%) (Fig. 2B). These isolates belonged to *Citrobacter*, *Enterobacter*, *Klebsiella*, and *Serratia*. The resistance mechanisms of such isolates to CFDC need to be analyzed in more detail in the future.

Among the tested *E. coli* isolates that were slightly less susceptible to CFDC (Fig. 4, MIC values between 1 and 4 µg/mL; the MIC values columns are shown in light blue), there were no isolates possessing all three CFDC-resistance factors (truncation of CirA, harboring a YRIN(K) insertion in PBP3, and harboring *bla*_NDM_). In other words, the combination of these three factors resulted in a stepwise decrease in susceptibility to CFDC. Seven *E. coli* isolates without these three CFDC-resistance factors showed slightly lower susceptibility to CFDC. This suggests the presence of other factors involved in the low susceptibility to CFDC. These results indicate that a combination of mechanisms contribute to reducing the susceptibility to CFDC, as previously described (30).

Two Enterobacterales isolates showing MIC values of 16 μg/mL against CFDC were susceptible to CAZ/AVI and IPM/REL (Fig. 3B). However, these strains did not possess any carbapenemases (data not shown). Thus, CAZ/AVI and IPM/REL show some potential against CFDC-resistant isolates. All five CFDC-resistant Enterobacterales strains were resistant to MEPM, DRPM, and IPM, suggesting that carbapenem use is not recommended for CFDC-resistant isolates. Fig. 3B, which shows the effective antibiotic agents against CFDC-resistant bacteria, could provide valuable information to combat CFDC-resistant bacteria in the clinical setting.

The antimicrobial efficacy of CFDC against IMP-type carbapenemase producers, which are dominant in Japan, is noteworthy (Fig. 2B). Further, KPC with D179Y (e.g., KPC-31 and −33) (31) or OXA-427 (32), known to confer resistance to CFDC, have not been found during surveillance in Japan (20). Therefore, CFDC should be categorized as a last resort in Japan; there are other effective classes of antimicrobial agents against CPEs in Japan (Fig. 3B). In addition, CAZ/AVI was effective against ESBL- or AmpC-harboring Enterobacterales (Fig. 3A). Appropriate use of antimicrobial agents in each clinical case is essential for saving the last resort, CFDC.

Among ESBL- or carbapenemase-producing and MEPM-nonsusceptible *E. coli* isolates obtained in our previous surveillance in Japan (JARBS-GNR) (19), there were eight isolates (0.25%) with both YRIN(K) insertion in PBP3 and truncated CirA (coverage ≦80%) including 3,158 strains other than the 71 strains used in this study. All eight strains belonged to ST167 or ST617 and seven of the eight strains possessed NDM-5 or NDM-9. In addition to the reports from China (17), *E. coli* ST167 isolated in Switzerland also showed a YRIN(K) insertion in PBP3 and truncated CirA (33). On this basis, it is presumed that potentially CFDC-resistant strains, even if they are currently susceptible, will not be missed in clinical settings. Because the proportion of these strains is extremely small, it is thought that the frequency of *E. coli* acquiring resistance to CFDC will be low in the future. This is considered an advantage of using CFDC in clinical settings in Japan.

A limitation of this study is the strains used in this study were selected based on strains obtained under specific conditions from previous surveillance, so the existence of potential bias cannot be ruled out. In this study, we examined clinical isolates in Japan where IMP-type MBLs are dominant and KPC type β-lactamases are rarely detected in CPE. CFDC showed robust in vitro efficacy against clinical isolates, including carbapenem nonsusceptible isolates.

## Conclusion

We focused on the effect of CFDC against a variety of CPEs, *P. aeruginosa*, and *Acinetobacter* spp., and carbapenemase non-producing Enterobacterales showing an MEPM MIC ≧2 μg/mL from clinical settings in Japan. This study enhances our understanding of the resistance mechanism against CFDC and helps establish the proper use of CFDC, CTLZ/TAZ, IPM/REL, CAZ/AVI, and other typical antibiotic agents in clinical settings.

## Data availability

The raw short- and long-read data were deposited in DDBJ under the BioProject accession number PRJDB10842 (https://www.ncbi.nlm.nih.gov/bioproject/?term=PRJDB10842), PRJDB16993 (https://www.ncbi.nlm.nih.gov/bioproject/?term=PRJDB16993), and PRJDB17296 (https://ddbj.nig.ac.jp/resource/bioproject/PRJDB17296).

## Acknowledgements

We thank Editage (www.editage.com) for the English language editing. We are also grateful to Sadao Aoki, Mikihisa Okuda, Hirokazu Yano, Akira Moriya, and Sayaka Uchino-Kondo for their technical contributions to this project. CFDC, CTLZ/TAZ, IPM/REL, and CAZ/AVI were kindly provided by the research laboratories of Shionogi & Co., Ltd. (Osaka, Japan). This study was supported by the Research Program on Emerging and Re-emerging Infectious Diseases of the Japanese Agency for Medical Research and Development (AMED) under grant numbers JP22fk0108604 and JP23fk0108604.

---

**Supplemental FIG 1** This figure shows the MIC_50/90_ values as a supplement to Fig. 2A. Unmeasured concentrations are diagonally lined (blue is shown as susceptible, yellow as intermediate resistance, and red as resistant). Dose-dependent sensitivity is indicated in green.

**Supplemental FIG 2** This figure shows the MIC_50/90_ values as a supplement to Fig. 2C and D. Unmeasured concentrations are diagonally lined (blue is shown as susceptible, yellow as intermediate resistance, and red as resistant).

